# A meiosis-specific factor C19orf57/4930432K21Rik/BRME1 modulates localization of RAD51 and DMC1 recombinases to DSBs in mouse meiotic recombination

**DOI:** 10.1101/2020.02.14.950204

**Authors:** Kazumasa Takemoto, Naoki Tani, Yuki Takada-Horisawa, Sayoko Fujimura, Nobuhiro Tanno, Mariko Yamane, Kaho Okamura, Michihiko Sugimoto, Kimi Araki, Kei-ichiro Ishiguro

## Abstract

Meiotic recombination is critical for genetic exchange and generation of chiasmata that ensures faithful chromosome segregation during meiosis I. Meiotic recombination is initiated by DNA double-strand break (DSB) followed by multiple processes of DNA repair. The exact mechanisms how recombinases localize to DSB remained elusive. Here we show that C19orf57/4930432K21Rik/BRME1 is a new player for meiotic recombination in mice. C19orf57/4930432K21Rik/BRME1 associates with ssDNA binding proteins, BRCA2 and MEILB2/HSF2BP, critical recruiters of recombinases onto DSB sites. Disruption of C19orf57/4930432K21Rik/BRME1 shows severe impact on DSB repair and male fertility. Remarkably, removal of single stranded DNA (ssDNA) binding proteins from DSB sites is delayed, and reciprocally the loading of RAD51 and DMC1 onto resected ssDNA is impaired in *Brme1* KO spermatocytes. We propose that C19orf57/4930432K21Rik/BRME1 modulates localization of recombinases to meiotic DSB sites through the interaction with the BRCA2-MEILB2/HSF2BP complex during meiotic recombination.

## Introduction

Meiosis consists of a single DNA replication followed by two rounds of chromosome segregation (meiosis I and meiosis II), which halves the chromosome number to ultimately produce haploid gametes. During meiotic prophase I, sister chromatids are organized into proteinaceous structures, termed axial element (AE) or chromosome axis (Zickler and Kleckner, 2015). Homologous chromosomes (homologs) then undergo synapsis, which is promoted by assembly of the synaptonemal complex (SC) (Cahoon and Hawley, 2016; Gerton and Hawley, 2005; Page and Hawley, 2004), and meiotic recombination yielding crossovers, a process that produces physical linkages between homologs called chiasmata.

Meiotic recombination is initiated by the introduction of DSB (Lam and Keeney, 2015) (Baudat et al., 2013) by SPO11 (Baudat et al., 2000; Romanienko and Camerini-Otero, 2000) and TOPO6BL (Robert et al., 2016; Vrielynck et al., 2016), and is completed by subsequent homologous recombination (HR)-mediated repair using homologs instead of sister chromatids as a template (Baudat and de Massy, 2007; Handel and Schimenti, 2010; Neale and Keeney, 2006). DNA-ends at DSBs are resected to form 3’-extended single stranded DNA (ssDNA) for invasion into a homologous template. ssDNA is coated by multiple ssDNA binding proteins, replication protein A (RPA) complexes (RPA1, RPA2, RPA3) (Wold et al., 1998) (Ribeiro et al., 2016), MEIOB (Luo et al., 2013) (Souquet et al., 2013), and SPATA22 (Ishishita et al., 2014; La Salle et al., 2012; Xu et al., 2017), to prevent degradation and secondary structure formation. Subsequently, RAD51 and DMC1 promote preceding removal of ssDNA binding proteins from ssDNA and facilitate invasion of 3’-extended strand into the duplex of the homolog (Cloud et al., 2012) (Kurzbauer et al., 2012) (Shinohara and Shinohara, 2004). In plants and mammals, BRCA2 directly interacts with RAD51 and DMC1, and plays an essential role in loading them onto ssDNA-ends (Jensen et al., 2010) (Sharan et al., 2004) (Siaud et al., 2004). Furthermore, recruitment of RAD51 and DMC1 on ssDNA is mediated by MEILB2/HSF2BP through the interaction with BRCA2 (Brandsma et al., 2019; Zhang et al., 2019). Although it has been proposed that MEILB2/HSF2BP-BRCA2 interaction plays a crucial role in the recruitment of RAD51 and DMC1 onto ssDNA, the exact mechanisms how the MEILB2/HSF2BP-BRCA2 complex acts for ssDNA processing remains elusive (Zhang et al., 2019).

Previously, we identified MEIOSIN that plays an essential role in meiotic initiation both in male and female (Ishiguro et al., 2020). MEIOSIN together with STRA8 acts as a crucial transcription factor to drive meiotic gene activation. *4930432K21Rik* gene was identified as one of the MEIOSIN/STRA8-regulated genes, whose expression was significantly downregulated in RNA-seq analysis of *Meiosin* KO (Ishiguro et al., 2020). And now, here we show that 4930432K21Rik, which we refer to as BRCA2 and MEILB2-associated protein1 (BRME1), is a novel meiotic recombination factor that modulates the loading of DMC1 and RAD51. We showed that BRME1 interacts with MEILB2/HSF2BP and BRCA2. Disruption of *Brme1* leads to defects in DSB repair and homolog synapsis, with severe impact on male fertility. In the absence of BRME1, removal of ssDNA binding proteins at the resected DNA-ends was delayed, and reciprocal recruitment of DMC1 and RAD51 was impaired during meiotic recombination. Present study suggests that BRME1 facilitates the recruitment of DMC1 and RAD51 to meiotic DSB sites through the interaction with the BRCA2-MEILB2/HSF2BP complex during meiotic recombination.

## Results

### C19orf57/BRME1 localizes along chromosome axis during meiotic prophase

A previously uncharacterized *4930432K21Rik* gene was identified as one of the MEIOSIN/STRA8-bound genes during preleptotene (FigS1A), whose expression was significantly downregulated in RNA-seq analysis of *Meiosin* KO (Ishiguro et al., 2020). Indeed, RT-PCR analysis demonstrated that *4930432K21Rik* expression level was down-regulated in *Meiosin* KO testis compared to postnatal day 10 (P10) wild type testes, where a cohort of spermatocytes undergo the first wave of meiotic entry (Fig. S1B).

We further examined the expression pattern of the *4930432K21Rik* in different mouse tissues using RT-PCR analysis. The *4930432K21Rik* gene showed a specific expression in adult testis and embryonic ovary but not in other adult organs that we examined (Fig. 1A, B), suggesting that *4930432K21Rik* is a germ-cell-specific factor. Public RNA-seq data of human cancer cells (Tang et al., 2017) indicated that human *4930432K21Rik* homolog was expressed not only in testis, but also in human tumors such as brain lower grade glioma, ovarian serous cystadenocarcinoma, and thymoma (Fig. S1C, see also Discussion). Mouse *4930432K21Rik* gene encodes a hypothetical protein C19orf57 homolog which possesses a conserved domain of unknown function (DUF4671). BLAST search analysis revealed that C19orf57 is conserved in vertebrates (Fig. S2). After its biological function described below, we referred to the protein product C19orf57 encoded by *4930432K21Rik* gene as the BRCA2 and MEILB2-associated protein1 (BRME1). To examine the subcellular localization patterns of this hypothetical protein, spread chromosomes of spermatocytes were immuno-stained with specific antibodies against BRME1 along with SYCP3 (a component of meiotic axial element: AE), and SYCP1 (a marker of homolog synapsis). The results showed that BRME1 protein appeared as foci along the chromosomes (Fig.1C). BRME1 foci faintly appeared on the nuclei at leptotene stage. Subsequently, the number of those foci culminated with more intense signals at zygotene, and declined from pachytene stage onward with residual foci persisting at diplotene. BRME1 foci were absent in *Spo11* KO spermatocytes (Fig.1D), whose DSB formation is defective (Lam and Keeney, 2015) (Baudat et al., 2013), indicating nuclear localization of BRME1 depends on DSBs. Notably, BRME1 foci were observed along unsynapsed axes, further implying its role at recombination nodules (Fig.1C). It should be mentioned that BRME1 foci showed less intense signals along synapsed chromosome, suggesting that BRME1 dissociated after homolog synapsis. Similarly, BRME1 protein appeared as foci in embryonic oocytes in a similar manner as spermatocytes, albeit faint signals (Fig.S4A). The observed spatiotemporal localization pattern of BRME1 protein resembled those of the factors involved in meiotic recombination, implying that BRME1 plays a role in the process of DSB repair during meiotic recombination.

**Figure 1.**
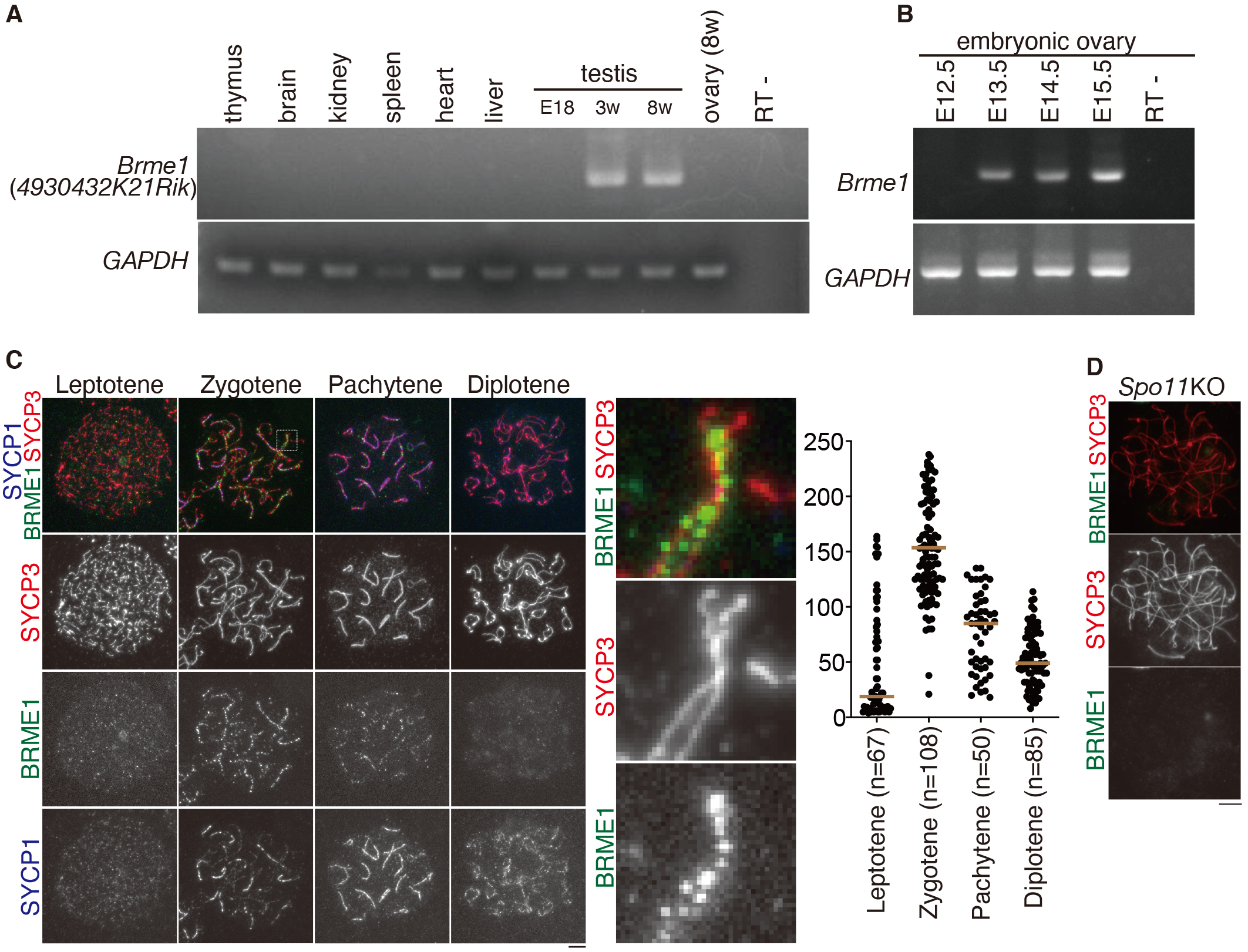
Identification of the novel meiosis-specific factor C19orf57/4930432K21Rik/BRME1. **(A)** The tissue-specific expression pattern of *4930432K21Rik/ Brme1* was examined using RT-PCR. Testis RNA was obtained from embryonic day 18 (E18.5) testis and tissues from adult eight-week-old male mice. RT-indicates control PCR without reverse transcription. (B) The expression pattern of *4930432K21Rik/ Brme1* in the embryonic ovary was examined using RT-PCR. Ovary RNA was obtained from E12.5-E15.5 female mice. **(C)** Chromosome spreads of WT spermatocytes were stained for BRME1, SYCP3, SYCP1. Scale bar: 5 μm. Enlarged images are shown to highlight axes that are going to be synapsed (middle). Numbers of BRME1 foci on SYCP3 axes are shown in the scatter plot with median (right). n indicates the number of cells examined. For immunostaining background, BRME1-immunostained signals were counted in *Brme1* KO. Note that numbers of BRME1 foci were significantly high in WT compared to those in *Brme1* KO. (see Fig. 2C) **(D)** Chromosome spreads of *Spo11* KO spermatocytes were stained for BRME1 and SYCP3. Scale bar: 5 μm.

### Disruption of *Brme1* led to severe defect in spermatogenesis

In order to address the role of *Brme1/4930432K21Rik* in meiosis, we deleted Exon3-Exon9 of *Brme1/4930432K21Rik* loci in C57BL/6 fertilized eggs through the CRISPR/Cas9 system (Fig. 2A). Western blotting of the extract from *Brme1* KO testis showed that BRME1 protein was absent (Fig. 2B), and immunolocalization of BRME1 along the chromosomes was diminished in *Brme1* KO (Fig. 2C), indicating that the targeted *Brme1/4930432K21Rik* allele was null. Although *Brme1* KO male mice did not show overt phenotype in somatic tissues, defects in male reproductive organs were evident with smaller-than-normal testes (Fig. 2D). Histological analysis revealed that the diameters of seminiferous tubules were reduced compared to those of WT or heterozygous mice, with slight residual presence of post-meiotic spermatids or sperms in a subpopulation of the tubules (59.8 % of total tubules) in eight-week-old *Brme1* KO testes (Fig. S3A). Close inspection of seminiferous tubules indicated that some, if any, seminiferous tubules of *Brme1* KO testis contained only a limited number of post-meiotic spermatids and sperms (Fig. S3A). Flow cytometry analysis of propidium-iodide (PI)-stained testicular cells isolated from 8-week old WT, *Brme1* +/- and *Brme1* KO mice showed that less population of 1N haploid cells were produced in *Brme1* KO mice compared to the control littermate (37.6% of 1N haploid cells in WT, 42.2% in *Brme1* +/- and 15.1 % in *Brme1* KO testicular cells) (Fig. S3B). Accordingly, the sperm density was largely reduced in *Brme1* KO caudal epididymis compared to those of the WT and *Brme1* +/- littermates (Fig. S3C). Mice heterozygous for the *Brme1* allele showed no obvious difference from WT (*Brme1*+/+) in fertility and histology in testes. To examine whether *Brme1* KO affected male fertility, mature *Brme1* KO males and their control littermates were mated with WT females over a period of 5-6 months. While the control males produced normal size of litters over this period, the *Brme1* KO males showed significantly reduced fertility (Fig. S3D). In contrast to male, *Brme1* KO females exhibited seemingly normal fertility with no apparent defect in adult ovaries (Fig. S3D, FigS4B).

**Figure 2.**
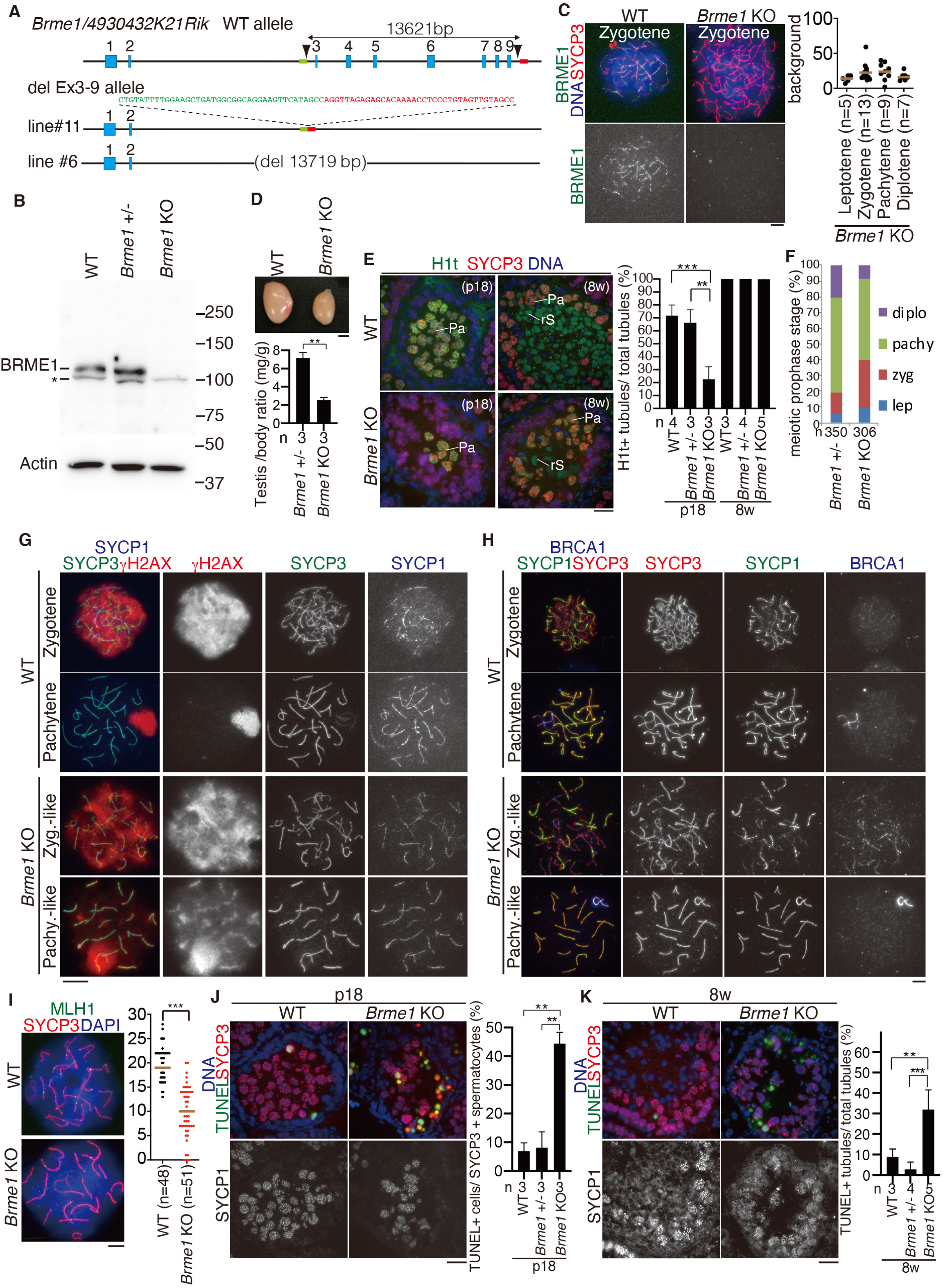
*Brme1* KO spermatocytes show defects in DSB repair and homolog synapsis. **(A)** The targeted Exon3-9 deletion allele of *Brme1* were generated by the introduction of CAS9, the synthetic gRNAs designed to target intron2 and the downstream of Exon9 (arrowheads), and ssODN into C57BL/6 fertilized eggs. Two lines of KO mice were established. Line #11 of *Brme1* KO mice was used in most of the experiments, unless otherwise stated. **(B)** Immunoblot analysis of testis extracts prepared from WT, *Brme1* +/- and *Brme1* KO mice (P18). Note that BRME1 protein migrated slower than the expected size calculated from the molecular weight. * indicates a non-specific band. **(C)** Chromosome spreads of WT and *Brme1* KO spermatocytes were stained for BRME1, SYCP3, and DAPI. Numbers of background BRME1-immunostained signals in *Brme1* KO axes are shown in the scatter plot with median (right). n indicates the number of cells examined. Note that the BRME1-immunostained signals in *Brme1* KO were considered to be background and the their numbers were negligibly small compared to those in WT as shown in Fig. 1C, confirming the specificity of the anti-BRME1 antibody. Scale bar: 5 μm. **(D)** Testes from WT, and *Brme1* KO mice (eight-week-old). Scale bar: 2 mm. Testis/body-weight ratio (mg/g) of *Brme1* +/- and *Brme1* KO mice (eight-week-old). n: number of animals examined. Statistical significance is shown by *p*-values (paired t-test). **: *p* <0.01 **(E)** Seminiferous tubule sections from WT and *Brme1* KO mice (P18, 8w) were stained as indicated. Pa: Pachytene spermatocyte, rS: round spermatid, eS: elongated spermatid. Shown below is the quantification of the seminiferous tubules that have H1t +/SYCP3+ cells per the seminiferous tubules that have SYCP3+ spermatocyte cells in WT (p18: n= 4, 8w: n=3), heterozygous (p18: n= 3, 8w: n=4) and *Brme1* KO (p18: n= 3, 8w: n=5) testes (bar graph with SD). n: the number of animals examined for each genotype. Statistical significance is shown by *p*-value (t-test). **: p < 0.01, ***: p < 0.001. Scale bar: 25 μm. **(F)** Chromosome spreads of *Brme1* +/- and *Brme1* KO spermatocytes (p20) were stained as indicated. Cells in the four developmental stages (leptotene, zygotene, pachytene and diplotene) were quantified in WT and *Brme1* KO. n: number of animals examined. lep: leptotene, zyg: zygotene, pachy: pachytene, diplo: diplotene. **(G)** Chromosome spreads of WT and *Brme1* KO spermatocytes were stained as indicated. **(H)** Chromosome spreads of WT and *Brme1* KO spermatocytes were stained as indicated. Zyg.-like: Zygotene-like, Pachy.-like: Pachytene-like. **(I)** Chromosome spreads of WT and *Brme1* KO spermatocytes were stained for MLH1, SYCP3 and DAPI. Statistical significance is shown by *p*-value (Mann-Whitney U-test). ***: p < 0.0001. Scale bars: 5 μm. **(J)** Seminiferous tubule sections from WT, *Brme1* +/- and *Brme1* KO mice (P18) were stained as indicated. Shown on the right is the quantification of the TUNEL+ cells per total SYCP3 + spermatocytes. Percentages of TUNEL+ cells were calculated in 16 seminiferous tubules, that have at least one TUNEL+ cell. Averaged percentage from 3 animals (total 48 tubules) for each genotype is shown with SD. Statistical significance is shown by *p*-values (paired t-test). **: *p* <0.01. Scale bar: 25 μm. **(K)** Seminiferous tubule sections from 8w mice were immunostained as in I. Shown on the right is the quantification of the seminiferous tubules that have TUNEL+ cells per total tubules in WT (8w: n= 3), *Brme1* +/- (8w: n=4) and *Brme1* KO (8w: n=5) testes (bar graph with SD). Statistical significance is shown by *p*-values (paired t-test). **: *p* <0.01. ***: p < 0.001. Scale bar: 25 μm.

We further investigated the fate of *Brme1* KO spermatocytes. Testis-specific histone H1t is a marker of spermatocytes later than mid pachytene. Immunostaining of seminiferous tubules by testis-specific histone H1t indicated that *Brme1* KO spermatocytes reached at least mid to late pachytene stage at P18, while the first wave of spermatogenesis had reached late pachytene stage in wild type (Fig. 2E). However, we noticed that the H1t positive population was markedly reduced in *Brme1* KO testes compared to WT and heterozygous controls at P18 (Fig. 2E), suggesting that progression of meiotic prophase was delayed or blocked in *Brme1* KO spermatocytes. Furthermore, counting the meiotic prophase population indicated that pachytene and diplotene populations were markedly reduced in *Brme1* KO spermatocytes (60.5 % versus 51.6 % in pachytene, 20.2 % versus 8.4 % in diplotene for control and *Brme1* KO, respectively) (Fig. 2F). Accordingly, zygotene population was apparently accumulated in *Brme1* KO spermatocytes (14.0 % versus 30.3 % for control and *Brme1* KO, respectively), suggesting that progression of meiotic prophase was compromised without BRME1. Therefore, although *Brme1* was not necessarily required for fertility in female, its absence had severe impact on male fertility.

### *Brme1* KO spermatocytes shows defects in DSB repair

DSB formation and repair are essential steps in meiotic recombination during meiotic prophase. Aforementioned results suggested that BRME1 protein acts for meiotic recombination (Fig. 1C). To determine the primary defect leading to subfertility in *Brme1* KO males, we analyzed the progression of meiotic prophase in *Brme1* KO testes. We examined DSB formation and repair events using immunostaining of γH2AX. A first wave of γH2AX is mediated by ATM after DSB formation at leptotene (Mahadevaiah et al., 2001), and disappears during DSB repair. The second wave of γH2AX at zygotene is mediated by ATR that targets unsynapsed chromosomes (Royo et al., 2013). At leptotene and zygotene, γH2AX signal appeared in *Brme1* KO spermatocytes in the same manner as WT (Fig. 2G), indicating that DSB formation normally occurred in *Brme1* KO spermatocytes. However, we noticed that γH2AX signals largely persisted throughout the nuclei until pachytene-like stage in *Brme1* KO spermatocytes, while they overall disappeared in WT pachytene spermatocytes except retaining on the XY body (Fig. 2G). This observation suggested that DSB repair was delayed or blocked in *Brme1* KO spermatocytes. Furthermore, BRCA1, a marker of asynapsis (Scully et al., 1997) (Broering et al., 2014), persisted along unsynapsed autosomal axes in zygotene-like *Brme1* KO spermatocytes (Fig. 2H), suggesting that meiotic silencing of unsynapsed chromatin (MUSC) was activated as a result of delayed homolog synapsis in *Brme1* KO spermatocytes. A subpopulation of *Brme1* KO spermatocytes indeed showed a pachytene morphology, whose 19 pairs of autosomal axes were apparently fully synapsed. However, the number of MLH1 foci, a marker of crossover (CO), was significantly reduced in *Brme1* KO pachytene-like spermatocytes compared to WT pachytene spermatocytes (Fig.2I). This implies that crossover recombination was incomplete or delayed in the absence of BRME1, despite the fact that a subpopulation of *Brme1* KO spermatocytes reached mid to late pachytene stage (Fig. 2E).

Consequently, TUNEL positive cells were robustly observed in the *Brme1* KO tubules (average 44.3 % of TUNEL positive spermatocytes in a given tubule) at P18, when the first wave of meiotic prophase reached late pachytene (Fig. 2J). The same phenotype persisted through adulthood in *Brme1* KO testes, because higher number of the TUNEL positive seminiferous tubules (∼31.9%) were observed in *Brme1* KO testis (Fig. 2K). TUNEL positive cells were exclusively observed in the seminiferous tubules that contained pachytene spermatocytes (Fig. 2J, K), and H1t positive population was reduced in *Brme1* KO testes compared to the controls at P18 (Fig. 2E). These observations suggested that a certain population of *Brme1* KO spermatocytes were consequently eliminated by apoptosis, probably leading to stage IV arrest. As a result, *Brme1* KO testes showed seminiferous tubules lacking post-meiotic spermatids or sperms at eight-week-old (Fig. S3A). Therefore, we reasoned that the primary defect in *Brme1* KO testes derived from failure of DSB repair during meiotic prophase.

### BRME1 facilitates recruitment of RAD51 and DMC1 recombinases to DSB sites

Given that BRME1 is required for DSB repair, we sought how BRME1 was involved in DSB repair processes by screening its interacting factors. Mass spectrometry (MS) analysis of immunoprecipitates of BRME1 from testes extracts identified MEILB2/HSF2BP (Brandsma et al., 2019; Yoshima et al., 1998; Zhang et al., 2019), BRCA2 (Sharan et al., 2004), RPA1(Wold et al., 1998) and MEIOB (Luo et al., 2013; Souquet et al., 2013), which are known to play a role in meiotic recombination (Fig.3A, Table S2). IP followed by western blotting using testes extracts indicated that BRME1 indeed coprecipitated those factors (Fig. 3B). This suggests that BRME1 mediates the process of meiotic recombination through the interaction with these factors. Notably, it was shown that MEILB2/HSF2BP directly interacts with BRCA2 and plays a role in recruiting recombinase RAD51 to DSBs through the interaction between BRCA2 and RAD51(Zhang et al., 2019). Indeed, BRME1 overall colocalizes with MEILB2/HSF2BP and MEIOB on the chromatin at zygotene (Fig.3C, D), while it rarely did with RAD51 and DMC1(Fig.3E, F).

**Figure 3.**
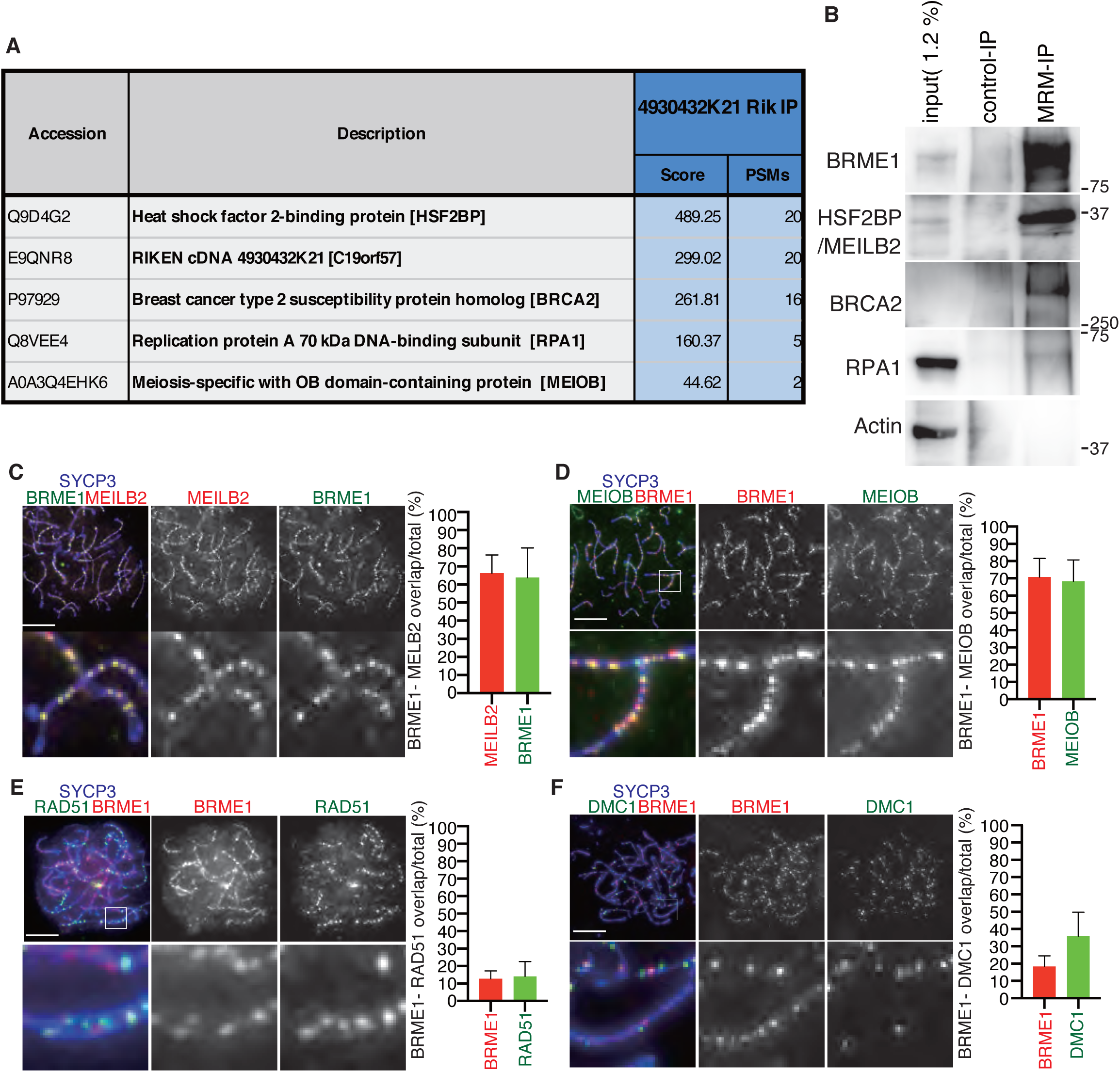
BRME1 interacts with BRCA2, MEILB2 and ssDNA binding proteins. **(A)** The immunoprecipitates of BRME1 from the chromatin-unbound fraction of the testis (P18) were subjected to liquid chromatography tandem-mass spectrometry (LC-MS/MS) analyses. The meiotic recombination proteins identified in the LC-MS/MS analyses of the samples are listed with the number of peptide hits and Mascot scores in the table. See table S2 for full list of raw MSMS data. **(B)** Immunoprecipitates of BRME1 from chromatin-unbound extracts of WT testes were immunoblotted as indicated. **(C-F)** Chromosome spreads of WT zygotene spermatocytes were stained as indicated. Scale bars: 5 μm. **(C)** Quantification of overlapped BRME1 and MEILB2 foci per total BRME1 or per total MEILB2 is shown (n= 10). **(D)** Quantification of overlapped BRME1 and MEIOB foci per total BRME1 or per total MEIOB is shown (n=11). **(E)** Quantification of overlapped BRME1 and RAD51 foci per total BRME1 or per total RAD51 is shown (n=15). **(F)** Quantification of overlapped BRME1 and DMC1 foci per total BRME1 or per total DMC1 is shown (n= 11).

We further monitored the dynamics of those factors involved in meiotic recombination in *Brme1* KO. Following DSB introduction, single strand DNA (ssDNA) binding proteins, RPA1, RPA2, RPA3 (Ribeiro et al., 2016), SPATA22 (Ishishita et al., 2014; La Salle et al., 2012; Xu et al., 2017) and MEIOB (Luo et al., 2013; Souquet et al., 2013), localize to the resected single stranded DNA-ends. Subsequently, DMC1 and RAD51 are recruited to the DNA-ends to promote strand invasion for homologous recombination (Pittman et al., 1998; Yoshida et al., 1998) (Cloud et al., 2012). In WT spermatocytes, ssDNA binding proteins, RPA2, MEIOB, and SPATA22 appeared at leptotene, culminated at zygotene, and declined toward pachytene as the resected ssDNA were repaired (Fig. 4A-C). In contrast, elevated number of RPA2 and SPATA22 foci were accumulated in *Brme1* KO spermatocytes compared to the controls (Fig. 4B-C), suggesting that the removal of RPA2 and SPATA22 was delayed or impaired in the absence of BRME1. We noticed that whereas elevated number of MEIOB foci were accumulated at zygotene in *Brme1* KO spermatocytes, comparable number of MEIOB foci were observed at pachytene in the control and *BRME1* KO spermatocytes (Fig. 4A). Since MEIOB rather persisted along the axes at pachytene irrespective of the presence or absence of BRME1 (Fig. 4A), MEIOB may have another role in the processing of joint molecules at later stage of recombination as proposed in the previous study (Luo et al., 2013).

**Figure 4.**
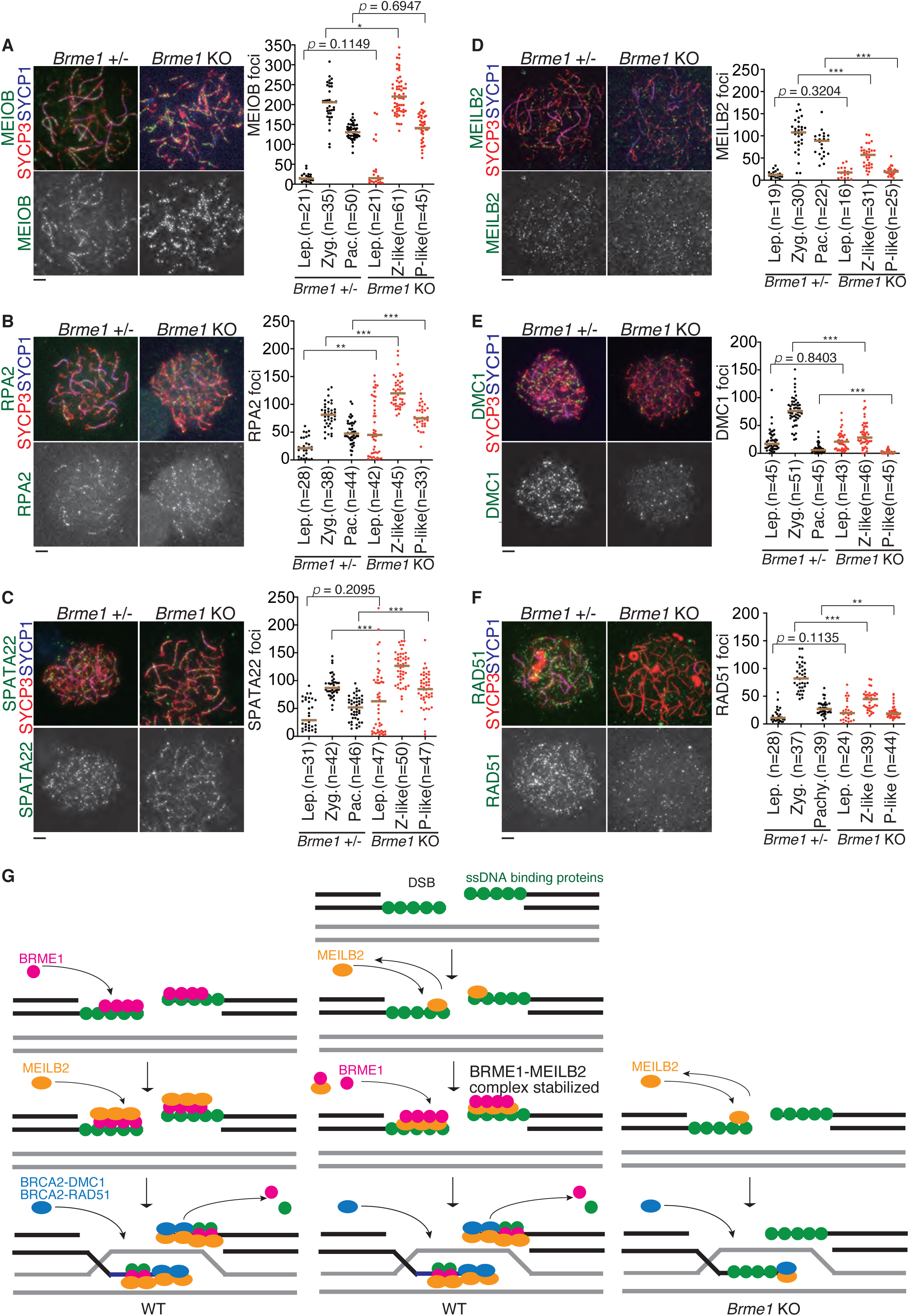
BRME1 plays a role in recruiting recombinases onto DSB sites. **(A-F)** Chromosome spreads of *Brme1* +/- and *Brme1* KO spermatocytes were stained as indicated. Immunostained chromosome spread of zygotene spermatocytes are shown. Scale bars: 5 μm. **(A)** The number of MEIOB foci is shown in the scatter plot with median (right). Statistical significance is shown by *p*-value (Mann-Whitney U-test). ***: p < 0.0001. **: p < 0.001. *: p < 0.05. Lep: leptotene, Zyg.: Zygotene, Pac.: Pachytene, Z-like: Zygotene-like, P-like: Pachytene-like. **(B)** The number of RPA2 foci is shown as in (A). **(C)** The number of SPATA22 foci is shown as in (A). **(D)** The number of MEILB2 foci is shown as in (A). **(E)** The number of DMC1 foci is shown as in (A). **(F)** The number of RAD51 foci is shown as in (A). **(G)** Schematic models of the role of BRME1 in meiotic recombination. Two alternative models are shown for WT (left, middle). Putative model is shown for *Brme1* KO (right). In *Brme1* KO, localization of MEILB2 onto DSBs are reduced. Consequently, the localization of recombinases is reduced. Black and grey lines indicate dsDNAs from homologous chromosomes.

Crucially, localization of MEILB2/HSF2BP was partly impaired in *Brme1* KO spermatocytes (Fig. 4D), suggesting that BRME1 facilitates localization of MEILB2/HSF2BP onto DSB sites or stabilizes MEILB2/HSF2BP on resected ssDNA. Accordingly, the number of RAD51 and DMC1 foci was significantly reduced in *Brme1* KO spermatocytes (Fig. 4E, F), suggesting that localization of RAD51 and DMC1 was partly destabilized or delayed in the absence of BRME1. However, *Brme1* KO showed less impact on localization of RAD51 and DMC1 than *Meilb2* KO, where RAD51 and DMC1 were completely abolished (Zhang et al., 2019). These results suggest that BRME1 at least in part facilitates recruitment of RAD51 and DMC1 recombinases to DSB sites through the interaction with MEILB2/HSF2BP and removes ssDNA binding proteins during meiotic recombination (Fig. 4G).

## Discussion

Present study revealed that BRME1/C19ORF57 plays a role in meiotic recombination. A key finding is that BRME1 interacts with MEILB2/HSF2BP. It has been shown that MEILB2/HSF2BP directly interacts with BRCA2 to promote loading of RAD51 and DMC1 recombinases onto DSB sites (Zhang et al., 2019). This process is mediated by two interactions: between BRCA2 and MEILB2/HSF2BP and between BRCA2 and RAD51/DMC1. However, how MEILB2/HSF2BP localizes to the DSB sites has been unclear. Present study demonstrates that BRME1 interacts with MEILB2/HSF2BP, BRCA2 and ssDNA binding proteins (Fig3A, B). Since armadillo repeats of MEILB2/HSF2BP directly interacts with BRCA2 (Brandsma et al., 2019; Zhang et al., 2019), BRCA2 was co-precipitated probably via MEILB2/HSF2BP in our BRME1 imunoprecipitates. BRME1 well colocalizes with MEILB2/HSF2BP and ssDNA binding proteins on the chromatin, while it rarely does with RAD51 and DMC1 (Fig3C-F). Furthermore, localizations of ssDNA binding proteins RPA, MEIOB and SPATA22 was elevated in *Brme1* KO spermatocytes, while those of MEILB2/HSF2BP, RAD51 and DMC1 were partly reduced (Fig. 4). These results suggest that BRME1 may initially bind to ssDNA-binding proteins and MEILB2/HSF2BP, then facilitate recruitment of the BRCA2-RAD51 and BRCA2-DMC1 complexes onto MEILB2/HSF2BP-bound DSB sites (Fig. 4G). Alternatively, MEILB2/HSF2BP associates with resected DSBs, and BRME1 subsequently interacts with MEILB2/HSF2BP leading to the stabilization of MEILB2/HSF2BP on DSBs (Fig. 5). It should be mentioned that removal of the ssDNA-binding protein RPA was severely delayed in *Brme1* KO (Fig. 4), whereas it was less affected in *Meilb*2 KO (Zhang et al., 2019), suggesting that BRME1 rather than MEILB2/HSF2BP may play more active role in replacing RPA from ssDNA. Thus, we speculate that BRME1 facilitates the loading of RAD51 and DMC1 recombinases onto DSBs through the interaction with MEILB2/HSF2BP and replacing ssDNA binding proteins.

We noticed that BRME1 persisted in late meiotic prophase (Fig 1C). Further, it was previously shown that MEILB2/HSF2BP persisted in pachytene (Zhang et al., 2019). It was demonstrated that early meiotic repair pathway that acts by default at the beginning of meiotic prophase is replaced sequentially by non-homologous end joining (NHEJ) and somatic-like homologous recombination (HR) pathway involving RAD51 (Enguita-Marruedo et al., 2019). Although we do not yet know the biological relevance of the detectable BRME1 foci in late meiotic prophase, BRME1 may be involved in “somatic-like HR repair pathway” in that stage.

In mouse, meiosis-specific ssDNA binding proteins MEIOB (Luo et al., 2013) (Souquet et al., 2013), and SPATA22 (Ishishita et al., 2014; La Salle et al., 2012; Xu et al., 2017) exist along with canonical ssDNA binding proteins RPA1-3. Thus, meiosis-specific ssDNA binding proteins have unique roles in processing homology-directed DNA repair during meiotic recombination. It has been shown that MEIOB interacts with SPATA22, and that their chromatin loading is interdependent (Luo et al., 2013). In the present study, we have shown that removal of SPATA22 was affected in *Brme1* KO spermatocytes (Fig. 4C), while that of MEIOB apparently was less affected (Fig. 4A). We do not know exact reason for different dynamics of SPATA22 and MEIOB in *Brme1* KO spermatocytes. However, this may be due to existence of another population of chromatin-bound SPATA22 and MEIOB at different time points, since it was previously proposed that MEIOB might mediate second-end capture after strand invasion (Luo et al., 2013).

DSS1 is widely conserved in vertebrates, nematode, plant and fungi (Kojic et al., 2003) (Marston et al., 1999) (Pispa et al., 2008) (Dray et al., 2006). In somatic cells, DSS1 interacts with BRCA2, and facilitates RPA-RAD51 exchange on ssDNA during homologous recombination (Zhao et al., 2015). Given that DSS1-BRCA2 interaction plays a role in replacing ssDNA binding protein with recombinases at DSBs, a similar mechanism may apply to BRME1-MEILB2/HSF2BP-BRCA2 interactions. As in the case of the previous study on MEILB2/HSF2BP (Zhang et al., 2019), requirement of BRME1 was also sexually dimorphic (Fig.S4). Although it is yet to be examined whether DSS1 or other factors compensates BRME1 in meiotic recombination, such redundant mechanisms may account for why residual sperm can be produced in *Brme1* KO males (Fig.S3), and BRME1 was not necessarily essential for female fertility (Fig. S4).

It should be mentioned that our *Brme1* KO male mice were subfertile, whereas those from the other group were completely infertile (Felipe-Medina et al. 2020). While our *Brme1* KO mice had larger deletion encompassing Exon3-Exon9 in the *Brme1* locus, which corresponded to almost the entire protein coding sequence, resulting in complete null allele (Fig.2A), those from the other group had 40bp deletion at the Exon 6, which was predicted to cause frame shift mutation. Although we do not know the exact reason for the phenotypic difference, it may be due to the difference in the alleles used in the two studies.

Lastly, it should be mentioned that expression levels of human *HSF2BP* is elevated in some tumors (Zhang et al., 2019) (Brandsma et al., 2019). Notably, expression levels of *BRME1/C19ORF57* was also elevated in the similar set of tumors (Fig. S1C) (Tang et al., 2017). Therefore, it is possible that both *BRME1/C19ORF57*and *MEILB2/HSF2BP* are co-expressed in a particular type of human tumors. This evidence supports the previously proposed hypothesis that ectopic expression of MEILB2 may perturb the intrinsic BRCA2 function through direct interaction in cancer cells (Zhang et al., 2019). Thus, misexpression of BRME1/C19ORF57 together with MEILB2 may compromise the function of the BRCA2-RAD51 complex during DNA repair in human cancers. Altogether, our study will shed light on multiple layers of mechanisms how recombinases are loaded onto DSB sites in vertebrate meiotic recombination.

## Acknowledgments

The authors thank Kumi Matsuura for technical support of ChIP-seq data reanalysis, Shoko Yamaoka for initial RT-PCR screening, Drs Satoshi Namekawa and Marry Ann Handel for provision of antibodies. This work was supported in part by KAKENHI in part by (#17H03634, #18K19304, #19H05245, #19H05743, #20H03265, #JP 16H06276) from MEXT, Japan; NIG-JOINT (32A2019); the program of the Joint Usage/ IMEG Research Center for Developmental Medicine; Takeda Science Foundation; Yamada Science Foundation; Ichiro Kanehara Foundation for Medical Science and Medical Care (to K.I.).

## Author contributions

K.T. performed the most of experiments. N.Tani performed MS analysis. Y.T. performed the RT-PCR and embryonic gonadal experiments. K.A. M.S. designed the knockout mice. N.Tanno performed flowcytometry. K.O. supported the antibody production. S.F. performed histological analyses. M.Y. performed bioinformatics analyses. K.I. supervised experiments, conducted the study and wrote the manuscript.

## STAR Methods

### Lead Contact and Materials Availability

Further information and requests for resources and reagents should be directed to and will be fulfilled by the Lead Contact, Kei-ichiro Ishiguro. *4930432K21Rik* knockout mouse lines generated in this study have been deposited to Center for Animal Resources and Development (CARD, ID 2775, 2776). The antibodies are available upon request. There are restrictions to the availability of antibodies due to the lack of an external centralized repository for its distribution and our need to maintain the stock. We are glad to share antibodies with reasonable compensation by requestor for its processing and shipping. All unique/stable reagents generated in this study are available from the Lead Contact with a completed Materials Transfer Agreement.

### Experimental Model and Subject Details

#### Animals

*4930432K21Rik/Brme1* knockout mice were C57BL/6 background. *Spo11* KO knockout mouse was reported earlier (Baudat et al., 2000). Male mice were used for immunoprecipitation of testis extracts, histological analysis of testes, immunostaining of testes, RT-PCR experiments. Female mice were used for histological analysis of the ovaries, immunostaining experiments. Whenever possible, each knockout animal was compared to littermates or age-matched non-littermates from the same colony, unless otherwise described. Animal experiments were approved by the Institutional Animal Care and Use Committee (approval F28-078, A2020-006, A30-001, A28-026).

### Method Details

#### Generation of *4930432K21Rik/Brme1* knockout mice and genotyping

*4930432K21Rik* knockout mouse was generated by introducing Cas9 protein (317-08441; NIPPON GENE, Toyama, Japan), tracrRNA (GE-002; FASMAC, Kanagawa, Japan), synthetic crRNA (FASMAC), and ssODN into C57BL/6N fertilized eggs using electroporation. For generating *4930432K21Rik/Brme1* Exon3-9 deletion (Ex3-9Δ) allele, the synthetic crRNAs were designed to direct GCAGGAAGTTCATAGCCACA(ggg) of the *4930432K21Rik* intron 2 and TGGGTAACAGATCTACACAC(agg) in the 3’-neighboring region of the Exon9. ssODN: 5’-GGCTACAACTACAGGGAGGTTTTGTGCTCTCTAACCTGTGaattcGGCTATGAACTTCCTGCCGCCATCAGCTTCCAAAATACAG-3’ was used as a homologous recombination template.

The electroporation solutions contained [10μM of tracrRNA, 10μM of synthetic crRNA, 0.1 μg/μl of Cas9 protein, ssODN (1μg/μl)] for *4930432K21Rik* knockout in Opti-MEM I Reduced Serum Medium (31985062; Thermo Fisher Scientific). Electroporation was carried out using the Super Electroporator NEPA 21 (NEPA GENE, Chiba, Japan) on Glass Microslides with round wire electrodes, 1.0 mm gap (45-0104; BTX, Holliston, MA). Four steps of square pulses were applied (1, three times of 3 mS poring pulses with 97 mS intervals at 30 V; 2, three times of 3 mS polarity-changed poring pulses with 97 mS intervals at 30 V; 3, five times of 50 mS transfer pulses with 50 mS intervals at 4 V with 40% decay of voltage per each pulse; 4, five times of 50 mS polarity-changed transfer pulses with 50 mS intervals at 4 V with 40% decay of voltage per each pulse).

The targeted *4930432K21Rik* Ex3-9Δ allele in F0 mice were identified by PCR using the following primers; 4930432K21Rik-1F: 5’-TGTTCACACAAAGTTCTTGAATCAG-3’ and 4930432K21Rik-3R: 5’-TCTGTGTGAGAATTGAGGCCTAAGC-3’ for the knockout allele (309 bp). 4930432K21Rik-2F: 5’-TAAATGATGCAGTCATGAGCCTCTG-3’ and 4930432K21Rik-1R: 5’-GGGGGTGATCAGAGCTCATTCCTAG-3’ for the Ex9 down stream of wild-type allele (345 bp). 4930432K21Rik-1F and 4930432K21Rik-2R: 5’-GGCTTTGAGGGATGGAGGCCGACCC-3’: for the intron 2 of wild-type allele (365 bp). The PCR amplicons were verified by sequencing. Primer sequences are listed in Table S1.

#### PCR with reverse transcription

Total RNA was isolated from tissues and embryonic gonads using TRIzol (Thermo Fisher). cDNA was generated from total RNA using Superscript III (Thermo Fisher) followed by PCR amplification using Ex-Taq polymerase (Takara) and template cDNA. For RT-qPCR, total RNA was isolated from WT, *Meiosin* KO and *Stra8* KO testes, and cDNA was generated as described previously (Ishiguro et al., 2020). *4930432K21Rik / Brme1* cDNA was quantified by ΔCT method using TB Green Premix Ex Taq II (Tli RNaseH Plus) and Thermal cycler Dice (Takara), and normalized by *GAPDH* expression level.

Sequences of primers used for RT-PCR (Fig1 A) are as follows:

GAPDH-F: 5’-TTCACCACCATGGAGAAGGC-3’

GAPDH-R: 5’-GGCATGGACTGTGGTCATGA-3’

4930432K21Rik-F1: 5’-GGCGACCTATTCCCCATCAG-3’

4930432K21Rik-R1: 5’-GGCCTTGTTTCCTGGGAAGG-3’

Sequences of primers used for RT-PCR (Fig1 B, Fig S1B) are as follows:

4930432K21Rik qPCR-F: 5’-AGTCACCAAACCTCAATCCA-3’

4930432K21Rik qPCR-R: 5’-AACCCCTTTGTCAGGTAAGG-3’

Gapdh_F2: 5’-ACCACAGTCCATGCCATCAC-3’

Gapdh_R2: 5’-TCCACCACCCTGTTGCTGTA-3’

Primer sequences are listed in Table S1.

#### Preparation of testis extracts and immunoprecipitation

Testis chromatin-bound and -unbound extracts were prepared as described previously (Ishiguro et al., 2014). Briefly, testicular cells were suspended in low salt extraction buffer (20 mM Tris-HCl [pH 7.5], 100 mM KCl, 0.4 mM EDTA, 0.1% TritonX100, 10% glycerol, 1 mM β-mercaptoethanol) supplemented with Complete Protease Inhibitor (Roche). After homogenization, the soluble chromatin-unbound fraction was separated after centrifugation at 100,000*g* for 10 min at 4°C. The chromatin bound fraction was extracted from the insoluble pellet by high salt extraction buffer (20 mM HEPES-KOH [pH 7.0], 400 mM KCl, 5 mM MgCl_2_, 0.1% Tween20, 10% glycerol, 1 mM β-mercaptoethanol) supplemented with Complete Protease Inhibitor. The solubilized chromatin fraction was collected after centrifugation at 100,000*g* for 10 min at 4°C.

For immunoprecipitation of endogenous 4930432K21Rik from chromatin unbound fraction, 3.75 µg of affinity-purified rat anti-4930432K21Rik and control IgG antibodies were crosslinked to 100 µl of protein A-Dynabeads (Thermo-Fisher) by DMP (Sigma). The antibody-crosslinked beads were added to the testis extracts prepared from wild type testes (17 to 21-day-old). The beads were washed with low salt extraction buffer. The bead-bound proteins were eluted with 40 µl of elution buffer (100 mM Glycine-HCl [pH 2.5], 150 mM NaCl), and then neutralized with 4 µl of 1 M Tris-HCl [pH 8.0]. The immunoprecipitated proteins were separately run on NuPAGE Bis-Tris 4%-12% gel (Thermo-Fisher) in MOPS-SDS buffer (Thermo-Fisher) for detection of BRME1, HSF2BP/MEILB, Actin, and on NuPAGE Tris-Acetate 3%-8% gel (Thermo-Fisher) in Tris-Acetate -SDS buffer (Thermo-Fisher) for detection of BRCA2, BRME1, HSF2BP/MEILB, RPA1, RAD51, DMC1, Actin. Immunoblots were detected by VeritBlot for IP detection reagent (HRP) (ab131366, abcam, 1:3000 dilution). Immunoblot image was developed using ECL prime (GE healthcare) and captured by FUSION Solo (VILBER). For the immunoblot of whole testes extracts from WT and *Brme1* KO mice, lysate were prepared in RIPA buffer and run on 8% Laemmli SDS-PAGE in Tris-Glycine-SDS buffer. Note that mobility of BRME1 protein was slower than expected molecular weight and slightly different depending on the gels and running buffers.

#### Mass spectrometry

The immunoprecipitated proteins were run on 4-12 % NuPAGE (Thermo Fisher) by 1 cm from the well and stained with SimplyBlue (Thermo Fisher) for the in-gel digestion. The gel containing proteins was excised, cut into approximately 1mm sized pieces. Proteins in the gel pieces were reduced with DTT (Thermo Fisher), alkylated with iodoacetamide (Thermo Fisher), and digested with trypsin and lysyl endopeptidase (Promega) in a buffer containing 40 mM ammonium bicarbonate, pH 8.0, overnight at 37°C. The resultant peptides were analyzed on an Advance UHPLC system (ABRME1ichrom Bioscience) coupled to a Q Exactive mass spectrometer (Thermo Fisher) processing the raw mass spectrum using Xcalibur (Thermo Fisher Scientific). The raw LC-MS/MS data was analyzed against the NCBI non-redundant protein/translated nucleotide database restricted to *Mus musculus* using Proteome Discoverer version 1.4 (Thermo Fisher) with the Mascot search engine version 2.5 (Matrix Science). A decoy database comprised of either randomized or reversed sequences in the target database was used for false discovery rate (FDR) estimation, and Percolator algorithm was used to evaluate false positives. Search results were filtered against 1% global FDR for high confidence level. All full lists of MSMS data are shown in Table S2 (Excel file).

#### Antibodies

The following antibodies were used for immunoblot (IB) and immunofluorescence (IF) studies: rabbit anti-Actin (IB, 1:1000, CST #4970), mouse anti-MLH1 (IF, 1:500, BD Biosciences: 551092), rabbit anti-SYCP1 (IF, 1:1000, Abcam ab15090), mouse anti-γH2AX (IF, 1:1000, Abcam ab26350), rabbit anti-DMC1 (IF, 1:500, Santa Cruz: SC-22768), rabbit anti-RAD51 (IF, 1:500, Santa Cruz: SC-8349), rabbit anti-SPATA22 (IF, 1:100, proteintech 16989-1-AP), rabbit anti-RPA1/RPA70 (IB, 1:1000, Abcam ab87272), rat anti-RPA2 (IF, 1:1000, CST 2208), rabbit anti-MEIOB (IF, 1:100, Millipore # ABE1414), mouse anti-SYCP1 (IF, 1:1000) (Ishiguro et al., 2011), rat anti-SYCP3 (Ishiguro et al., 2020) (IF, 1:1000), gunia pig anti-SYCP3 (Ishiguro et al., 2020) (IF, 1:2000), goat anti-BRCA1 (IF, 1:500, SantaCruz sc-1553), rabbit anti-BRCA2 (IB, 1:1000, Abcam ab27976), guinea pig anti-H1t (IF, 1:2000, kindly provided by Marry Ann Handel).

Polyclonal antibodies against mouse BRME1/4930432K21Rik-N and mouse MEILB2/HSF2BP were generated by immunizing rabbits and rats. His-tagged recombinant proteins of BRME1/4930432K21Rik-N (aa 1-207) and MEILB2/HSF2BP (Full length) were produced by inserting cDNA fragments in-frame with pET28c (Novagen) in *E. coli* strain BL21-CodonPlus(DE3)-RIPL (Agilent), solubilized in a denaturing buffer (6 M HCl-Guanidine, 20 mM Tris-HCl [pH 7.5]) and purified by Ni-NTA (QIAGEN) under denaturing conditions. The antibodies were affinity-purified from the immunized serum with immobilized antigen peptides on CNBr-activated Sepharose (GE healthcare). Rat MEILB2/HSF2BP antibody was labeled by Alexa555 using Alexa Fluor 555 Antibody labeling kit (Thermo A20187).

#### Histological Analysis

Testes, caudal epididymis and ovaries were fixed in Bouin’s solution, and embedded in paraffin. Sections were prepared on CREST-coated slides (Matsunami) at 6 μm thickness. The slides were dehydrated and stained with hematoxylin and eosin. For Immunofluorescence staining, testes were embedded in Tissue-Tek O.C.T. compound (Sakura Finetek) and frozen. Cryosections were prepared on the CREST-coated slides (Matsunami) at 8 μm thickness, and then air-dried. The serial sections of frozen testes were fixed in 4% paraformaldehyde in PBS for 5 min at room temperature and washed briefly in PBS. After washing, the serial sections were permeabilized in 0.1% TritonX100 in PBS for 5 min. The sections were blocked in 3% BSA/PBS, and incubated at room temperature with the primary antibodies in a blocking solution. After three washes in PBS, the sections were incubated for 1 h at room temperature with Alexa-dye-conjugated secondary antibodies (1:1500; Invitrogen) in a blocking solution. TUNEL assay was performed using MEBSTAIN Apoptosis TUNEL Kit Direct (MBL 8445). DNA was counterstained with Vectashield mounting medium containing DAPI (Vector Laboratory).

#### Immunostaining of spermatocytes

Surface-spread nuclei from spermatocytes and oocytes were prepared by the dry down method as described (Peters et al., 1997) with a modification. The sides were then air-dried and washed with water containing 0.1 % Tween20 or frozen for longer storage at - 30°C. For staining of HSF2BP/MEILB2, the slides were heat-treated briefly by microwave oven in TE (pH8.0) for antigen retrieval. The slides were permeabilized in 0.1% TritonX100 in PBS for 5 min, blocked in 3% BSA/PBS, and incubated at room temperature with the primary antibodies in 3% BSA/PBS. For immunostaining of SPATA22, slides were blocked in PBS containing 5% skim milk and 5% FBS. After three washes in PBS, the sections were incubated for 1 h at room temperature with Alexa-dye-conjugated secondary antibodies (1: 1500; Invitrogen) in a blocking solution. DNA was counterstained with Vectashield mounting medium containing DAPI (Vector Laboratory).

#### Imaging

Immunostaining images were captured with DeltaVision (GE Healthcare). The projection of the images was processed with the SoftWorx software program (GE Healthcare). All images shown were Z-stacked. Bright field images were captured with OLYMPUS BX53 fluorescence microscope and processed with CellSens standard program. For counting seminiferous tubules, immunostaining images were captured with BIOREVO BZ-X710(KEYENCE), and processed with BZ-H3A program.

#### FACS analysis

Testes were scraped and digested with accutase (Innovative cell technologies Inc.) at room temperature in the presence of DNase II followed by filtration through a 40 μm cell strainer (FALCON). The testicular cells were fixed in 70% ethanol for over night, washed twice with 1 % BSA in PBS and brought to a concentration of 1 × 10^6^ cells/ml in propidium iodide/RNase solution (50μg/mL). DNA content was analyzed using SP6800 spectral analyzer (Sony Biotechnology Inc., Tokyo, Japan) with 488nm laser illumination. Data were analyzed and processed using FlowJo for PC, version 10.0.7 (Becton, Dickinson & Company, Ashland, OR).

#### ChIP-seq Data and Public RNA-seq data Analysis

MEIOSIN and STRA8 ChIP-seq data were described in our previous study (Ishiguro et al., 2020), and analyzed for *4930432K21Rik* / *Brme1* locus. MEIOSIN and STRA8 binding sites were shown along with genomic loci from Ensembl on the genome browser IGV.

Public RNA-seq data was analyzed using GEPIA2 server http://gepia2.cancer-pku.cn/#index (Tang et al., 2017) at the threshold of Log2FC Cutoff:1, p-value Cutoff:0.01.

#### Quantification and Statistical analysis

Statistical analysis was performed using GraphPad Prism or Microsoft EXCEL.

**Figure 1A** RT-PCR data was taken from an adult male (1 animal) and an adult female (1 animal).

**Figure 1B** RT-PCR data was taken from female embryos at indicated time points (one embryo for each time point).

**Figure 1C** Foci counting data was pooled from 3 males.

**Figure 1D** Data was acquired from *Spo11* KO male (1 animal).

**Figure 2B** BRME1 protein level was examined by western blotting from WT (P18; 2 animals), *Brme1* +/- (P18; 1 animal) and *Brme1* KO (P18; 2 animals).

**Figure 2C** Background signals of BRME1 immunostaining were counted at each stage from WT (1 animal) and *Brme1* KO (1 animal). n: the number of cells examined at each stage.

**Figure 2D** Quantification of a pair of testes/body-weight ratio (mg/g) in *Brme1* +/- (8w; n=3) and *Brme1* KO (8w; n=3) mice. n: the number of animals examined for each genotype. Bar graph indicates mean with SD. Statistical significance was determined by t-test.

**Figure 2E** Quantification of the seminiferous tubules that have H1t +/SYCP3+ cells per the seminiferous tubules that have SYCP3+ spermatocyte cells in WT (p18: n= 4, 8w: n=3), heterozygous (p18: n= 3, 8w: n=4) and *Brme1* KO (p18: n= 3, 8w: n=5) testes. n: the number of animals examined for each genotype. Bar graph indicates mean with SD. Statistical significance was determined by t-test.

**Figure 2F** Spermatocytes in the four developmental stages (leptotene, zygotene, pachytene, and diplotene) per total cells in meiotic prophase were quantified in WT (n=350 from one animal) and *Brme1* KO (n=306 from one animal).

**Figure 2I** Numbers of MLH1 foci on SYCP3 axes were counted in WT and *Brme1* KO. Number of foci was indicated in the scatter plot with median. *p*-value (Mann-Whitney U-test) is shown.

**Figure 2J** Quantification of the TUNEL+ cells per total SYCP3 + spermatocytes in WT, *Brme1* +/- and *Brme1* KO mice (P18). Percentages of TUNEL+ cells per total SYCP3 + spermatocytes were calculated in 16 seminiferous tubules, that have at least one TUNEL+ cell. Averaged percentage from 3 animals (total 48 tubules) for each genotype is calculated. Bar graph indicates mean with SD. Statistical significance was determined by t-test.

**Figure 2K** Quantification of the seminiferous tubules that have TUNEL+ cells per total tubules in WT (8w: n=3), *Brme1* +/- (8w: n=4) and *Brme1* KO (8w: n=5) testes. Bar graph indicates mean with SD. Statistical significance was determined by t-test.

**Figure 3C** Quantification of overlapped BRME1 and MEILB2 foci per total BRME1 or per total MEILB2 is shown (n= 10). Bar graph indicates mean with SD.

**Figure 3D** Quantification of overlapped BRME1 and MEIOB foci per total BRME1 or per total MEIOB is shown (n=11). Bar graph indicates mean with SD.

**Figure 3E** Quantification of overlapped BRME1 and RDA51 foci per total BRME1 or per total RDA51 is shown (n=15). Bar graph indicates mean with SD.

**Figure 3F** Quantification of overlapped BRME1 and DMC1 foci per total BRME1 or per total DMC1 is shown (n= 11). Bar graph indicates mean with SD.

**Figure 4A-F** Numbers of MEIOB, RPA2, SPATA22, MEILB2, DMC1 and RAD51 foci on SYCP3 axes were counted in WT and *Brme1* KO. Number of foci was indicated in the scatter plot with median. *p*-value (Mann-Whitney U-test) is shown.

**Figure S1B** Testis RNA was obtained from P8 WT (3 animals), P10 WT (4 animals), *Meiosin* KO (3 animals) and *Stra8* KO (4 animals). qPCR was performed in triplicates, and the average ddCt values were calculated for individual cDNA samples. The expression level of *Brme1* was divided by that of *GAPDH* to give a relative expression level of *Brme1* to *GAPDH.* Relative expression level of *Brme1* to *GAPDH* was normalized to 1 for a given P10 WT sample. Bar graph indicates mean with SD. Statistical significance was determined by t-test.

**Figure S1C** The box plot was generated using GEPIA server (Tang et al., 2017). Red box indicates tumor. Grey box indicates normal tissue. The numbers of normal (num(N)) and tumor (num(T)) samples are indicated. Statistical significance was determined by one-way ANOVA.

**Figure S3A** Quantification of the seminiferous tubules containing post-meiotic cells per total number of the seminiferous tubules in WT (n=3), *Brme1* +/- (n=4) and *Brme1* KO (n=5) mice. Bar graph indicates mean with SD. Statistical significance was determined by t-test.

**Figure S3B** Quantification of 1N post-meiotic cells from flow cytometry histogram were performed using Flowjo. Bar graph indicates mean with SD. Data was acquired from WT (8w; 3 animals), *Brme1* +/- (8w; 3 animals) and *Brme1* KO (8w; 3 animals). Statistical significance was determined by t-test.

**Figure S4A** Data was acquired from WT (1animal) and *Brme1* KO female embryos (1 animal).

**Figure S4B** Adult ovaries examined by HE staining were acquired from *Brme1* +/- (1 animal) and *Brme1* KO (2 animals) females.

## DATA AND CODE AVAILABILITY

All data supporting the conclusions are present in the paper and the supplementary materials. Original data are deposited in Mendeley Data (http://dx.doi.org/10.17632/jjmg95zkyx.1). Sequencing data for the mouse homolog of *C19orf57/4930432K21Rik* gene was initially deposited in the National Center for Biotechnology Information-National Institutes of Health (NCBI-NIH) GenBank under accession numbers LC507101 as *Miotic recombination modulator* (*Mrm*), and posted on preprint server bioRxiv (https://doi.org/10.1101/2020.02.14.950204). Later, we accepted the nomenclature *Brme1* that was given to *4930432K21Rik*. Mouse lines generated in this study have been deposited to Center for Animal Resources and Development (CARD), *4930432K21Rik* mutant mouse #11 (ID 2775) and *4930432K21Rik* mutant mouse #6 (ID 2776).

## Supplemental items

**Table S1 (Excel file)**

Primers and oligos used in this study, related to STAR Methods and the Key Resources Table

**Table S2 (Excel file)**

Full list of peptides identified in control IgG IP and 4930432K21rik IP The following data are shown in the tabs.

(1) Full list of peptides identified in control IgG IP and 4930432K21rik IP are shown.

(2) 4930432K21rik IP specific peptides are shown after subtraction of the peptides identified in control IgG IP and IgG fragments.

(3) The images of gel slices used for shotgun LC-MSMS are shown.

(4) Raw MSMS data for the peptides identified in control IgG IP

(5) Raw MSMS data for the peptides identified in 4930432K21rik IP

**Supplementary Figure 1.**
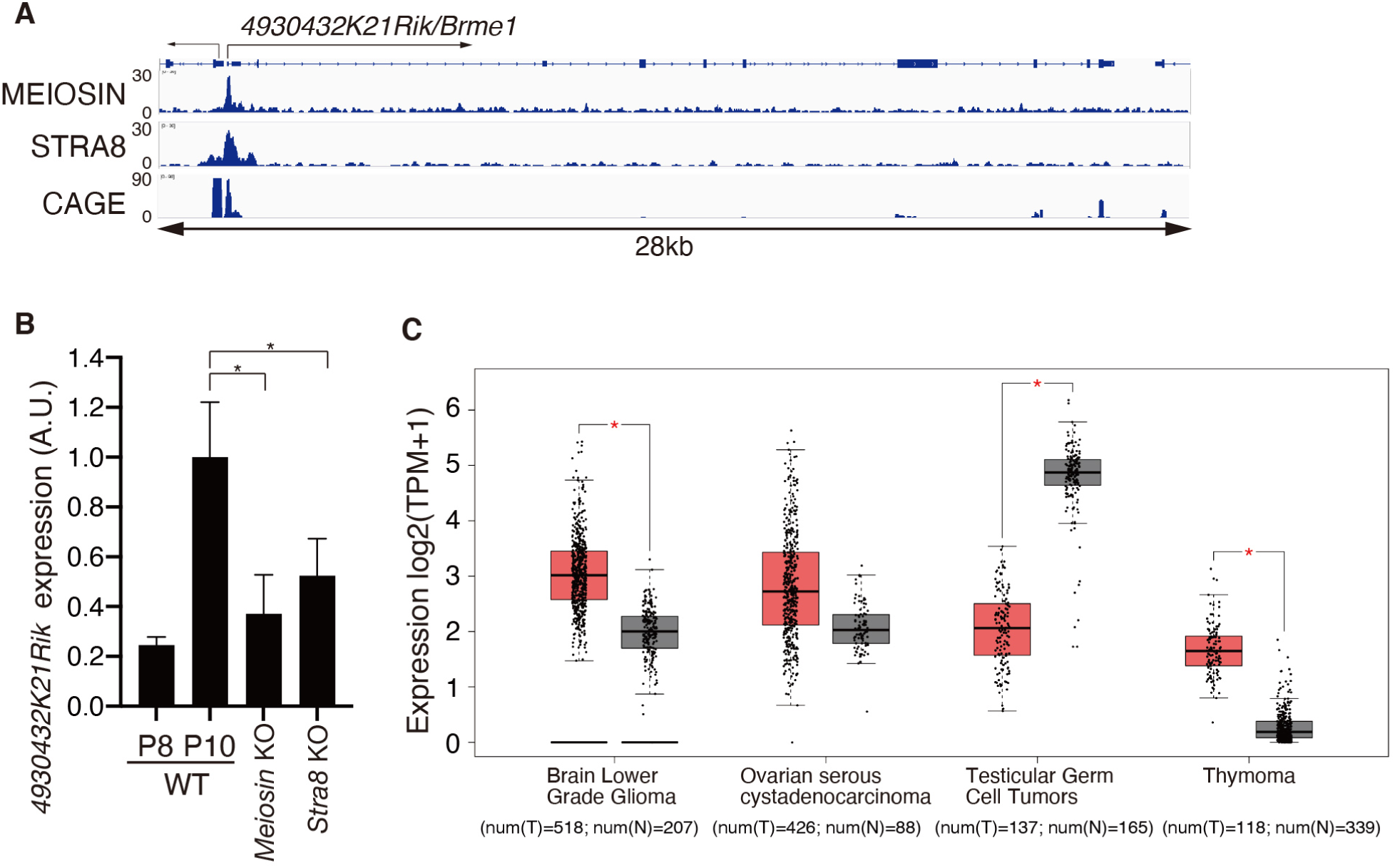
Identification of the meiosis-specific factor 4930432K21 Rik /C19orf57/ BRME1 (related to Figure 1) **(A)** Genomic view of MEIOSIN and STRA8 binding peaks over *4930432K21Rik/Brme1* loci. To specify testis specific TSS, RNA-seq of the 5’ capped end of the mRNA (CAGE) in P10.5 testis is shown (41). Genomic coordinates were obtained from Ensembl. **(B)** The expression of *4930432K21Riklbrme1* in WT versus Meiosin KO was examined using RT-PCR. Testis RNA was obtained from WT (P8 and P10), *Meiosin* KO and *Stra8* KO. *p*-value (t-test) is shown by statistical significance. *: p < 0.01. **(C)** Expression of *MRMIC19orf57* in human tumors and normal tissues of different origin. The plot was generated from public RNA-seq data using GEPIA2 server (Tang et al., 2017). Red box indicates tumor. Grey box indicates normal tissue. Number of normal (num(N)) and tumor (num(T)) samples is indicated; Tpm – transcripts per kilobase million. Statistical significance was determined by one-way ANOVA.

**Supplementary Figure 2.**
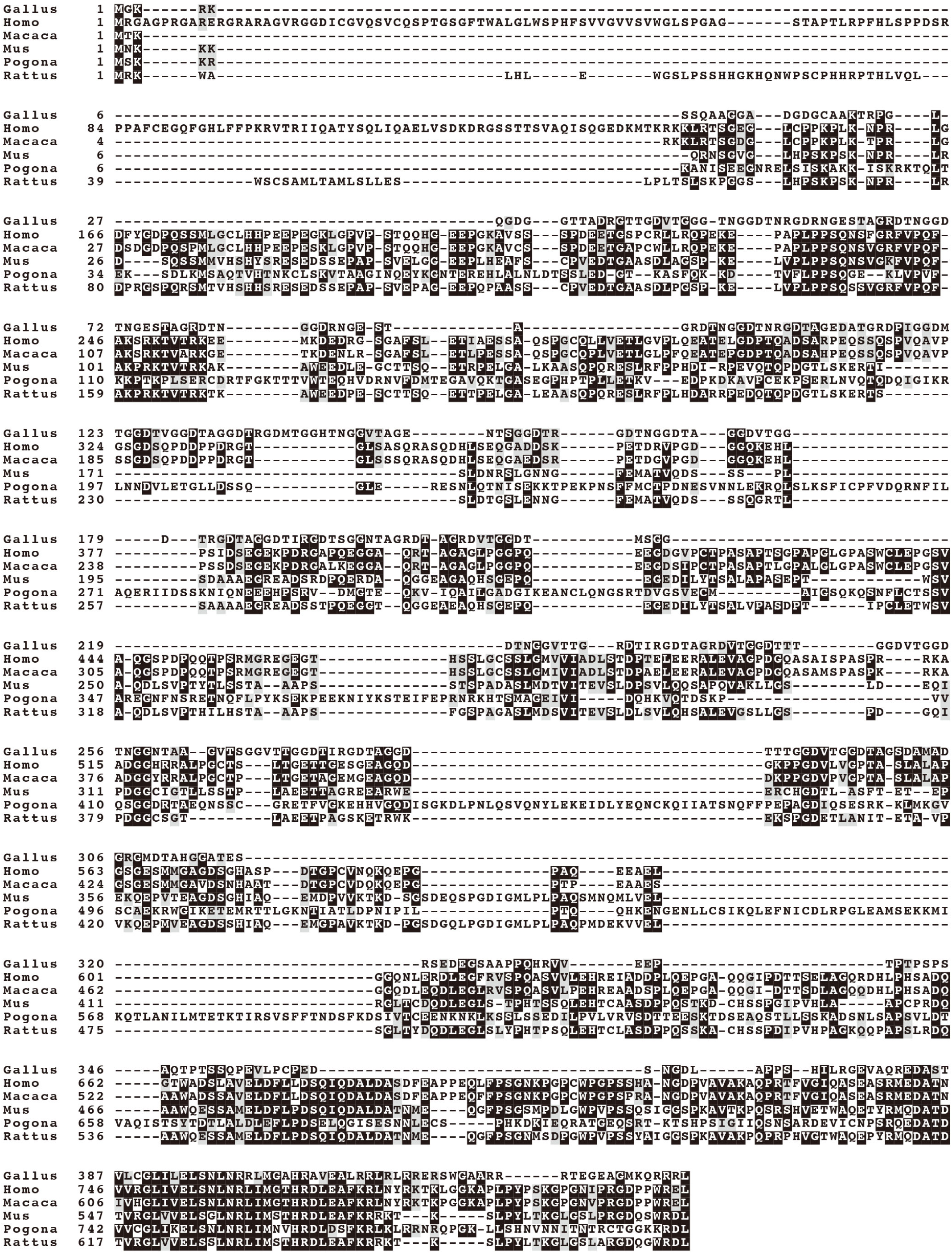
Sequence alignment of BRME1/C19orf57 homologs in vertebrates (related to Figure 1) BRME1/C19orf57 protein is conserved among vertebrates. The amino acid sequence of *M. musculus* BRME1/C19orf57/MRM (mouse, LC507101) derived from this study. C19orf57 homologs of *R. norvegicus* (rat, XP _008770650.1), *H. sapiens* (human, NP _001332772.1), *Macaca mulatta* (monkey, XP_001111192.4), *Pogona vitticeps* (reptile, XP_020654389.1) and *Gallus gal/us* (chick, XP_025001559.1) were obtained from the National Center for Biotechnology Information-National Institutes of Health (NCBI-NIH) protein database.

**Supplementary Figure 3.**
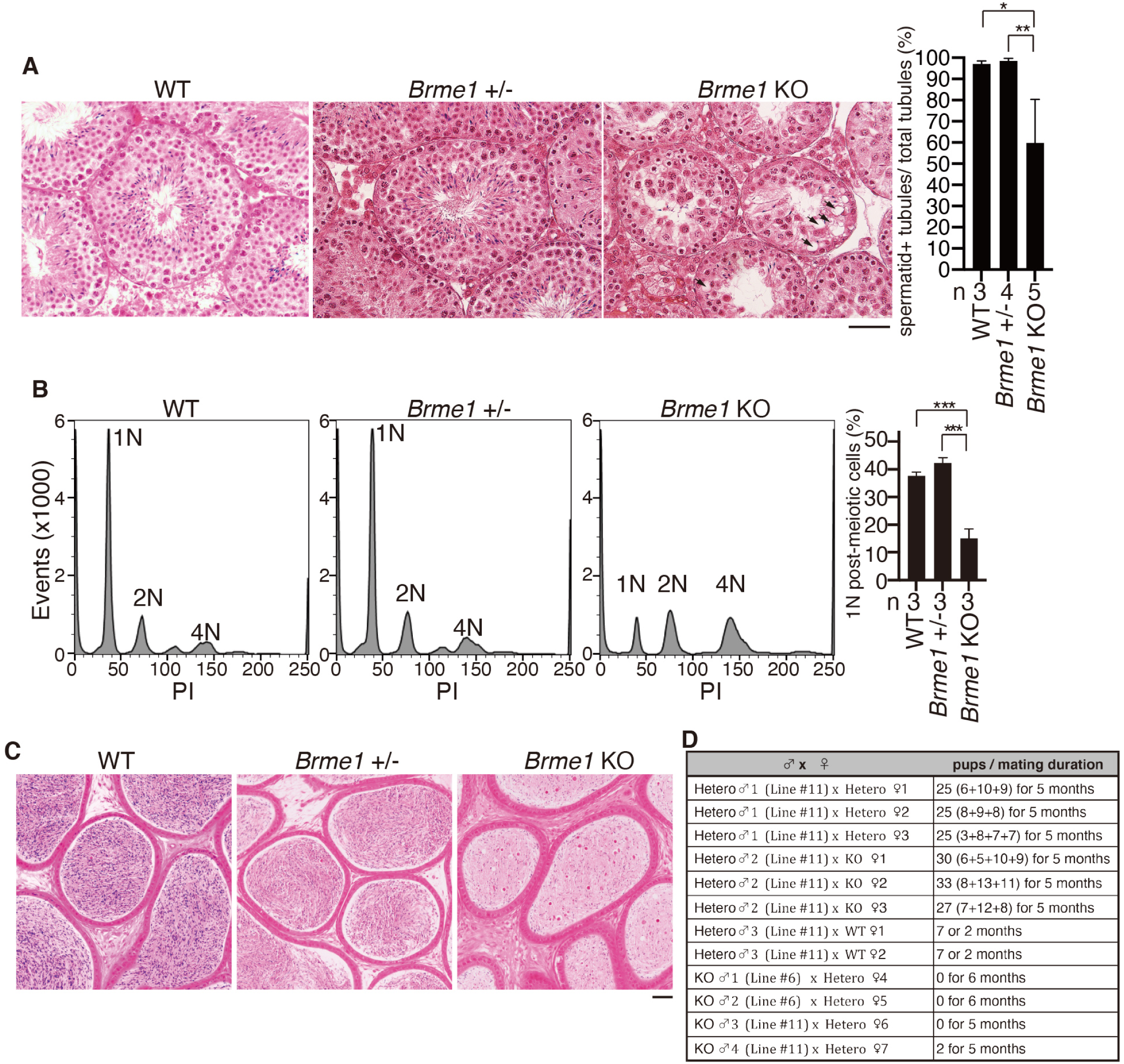
Spermatogenesis was impaired in *Brme1* KO testis (related to Figure 2) **(A)** Hematoxylin and eosin staining of the sections from WT, *Brme1* +/- and *Brme1* KO testes (eight-week-old). Arrow indicates a vacuole. Scale bar: 50 µm. Quantification of the seminiferous tubules containing post-meiotic cells per total number of the seminiferous tubules in WT testes (bar graph with SD). n: the number of the animals examined for each genotype. *p*-values are shown (paired t-test). **: *p* <0.01. *: *p* < 0.05. **(B)** Flow cytometry histogram of propidium-iodide (Pl)-stained testicular cells isolated from 8-week old *Brme 1* +/+ (WT), *Brme1* +/- and *Brme1* KO mice. N indicates DNA content in the cell population. Quantification of 1N post-meiotic cells is shown (bar graph with SD, n=3 for each genotype). *p*-values are shown (paired t-test). ***: *p* <0.001. **(C)** Hematoxylin and eosin staining of the sections from *Brme1* +/+(WT), *Brme1* +/- and *Brme1* KO epididymis (eight-week-old). Scale bar: 50 µm. **(D)** Fertility of *Brme1* KO males and females was examined by mating for the indicated period. Note that apparent differences were not observed between *Brme 1* +/+(WT) and *Brme 1* +/- testes.

**Supplementary Figure 4.**
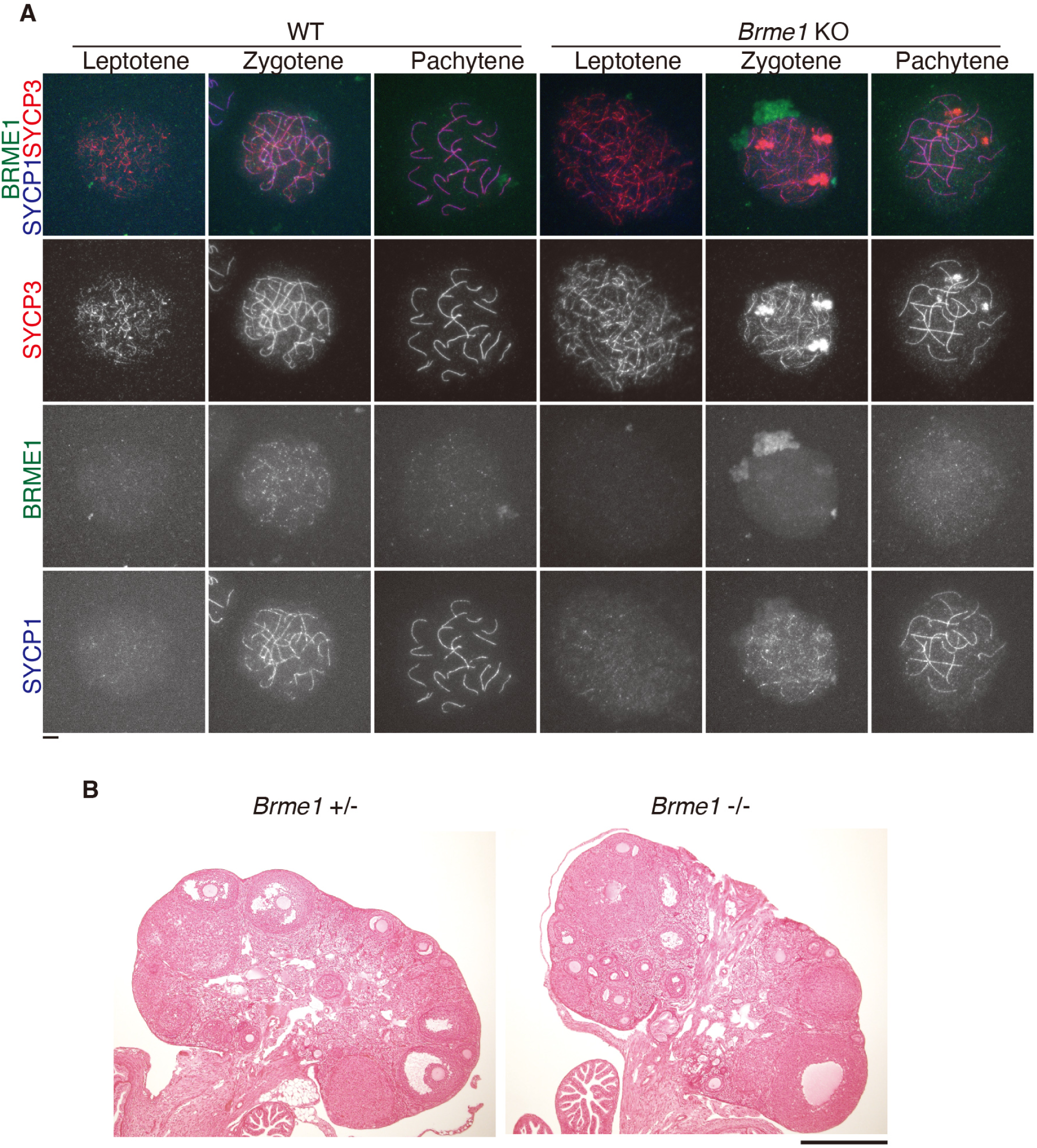
Phenotypic analyses of *Brme1* KO ovaries (related to Figure 2) **(A)** Chromosome spreads of oocytes in WT and *Brme1* KO (KO allele #6) were stained for BRME1, SYCP3, and SYCP1. Scale bars: 5 µm. **(B)** Hematoxylin and eosin staining of the sections from *Brme1* +/- and *Brme1* KO (KO allele #6) ovaries (eight-week-old). Scale bar: 500 µm.

## References

Baudat, F., and de Massy, B. (2007). Regulating double-stranded DNA break repair towards crossover or non-crossover during mammalian meiosis. Chromosome Res 15, 565–577.

Baudat, F., Imai, Y., and de Massy, B. (2013). Meiotic recombination in mammals: localization and regulation. Nat Rev Genet 14, 794–806.

Baudat, F., Manova, K., Yuen, J.P., Jasin, M., and Keeney, S. (2000). Chromosome synapsis defects and sexually dimorphic meiotic progression in mice lacking Spo11. Mol Cell 6, 989–998.

Brandsma, I., Sato, K., van Rossum-Fikkert, S.E., van Vliet, N., Sleddens, E., Reuter, M., Odijk, H., van den Tempel, N., Dekkers, D.H.W., Bezstarosti, K., et al. (2019). HSF2BP Interacts with a Conserved Domain of BRCA2 and Is Required for Mouse Spermatogenesis. Cell Rep 27, 3790–3798 e3797.

Broering, T.J., Alavattam, K.G., Sadreyev, R.I., Ichijima, Y., Kato, Y., Hasegawa, K., Camerini-Otero, R.D., Lee, J.T., Andreassen, P.R., and Namekawa, S.H. (2014). BRCA1 establishes DNA damage signaling and pericentric heterochromatin of the X chromosome in male meiosis. J Cell Biol 205, 663–675.

Cahoon, C.K., and Hawley, R.S. (2016). Regulating the construction and demolition of the synaptonemal complex. Nat Struct Mol Biol 23, 369–377.

Cloud, V., Chan, Y.L., Grubb, J., Budke, B., and Bishop, D.K. (2012). Rad51 is an accessory factor for Dmc1-mediated joint molecule formation during meiosis. Science 337, 1222–1225.

Dray, E., Siaud, N., Dubois, E., and Doutriaux, M.P. (2006). Interaction between Arabidopsis Brca2 and its partners Rad51, Dmc1, and Dss1. Plant Physiol 140, 1059–1069.

Enguita-Marruedo, A., Martin-Ruiz, M., Garcia, E., Gil-Fernandez, A., Parra, M.T., Viera, A., Rufas, J.S., and Page, J. (2019). Transition from a meiotic to a somatic-like DNA damage response during the pachytene stage in mouse meiosis. PLoS Genet 15, e1007439.

Gerton, J.L., and Hawley, R.S. (2005). Homologous chromosome interactions in meiosis: diversity amidst conservation. Nat Rev Genet 6, 477–487.

Handel, M.A., and Schimenti, J.C. (2010). Genetics of mammalian meiosis: regulation, dynamics and impact on fertility. Nat Rev Genet 11, 124–136.

Ishiguro, K., Kim, J., Fujiyama-Nakamura, S., Kato, S., and Watanabe, Y. (2011). A new meiosis-specific cohesin complex implicated in the cohesin code for homologous pairing. EMBO Rep 12, 267–275.

Ishiguro, K., Kim, J., Shibuya, H., Hernandez-Hernandez, A., Suzuki, A., Fukagawa, T., Shioi, G., Kiyonari, H., Li, X.C., Schimenti, J., et al. (2014). Meiosis-specific cohesin mediates homolog recognition in mouse spermatocytes. Genes Dev 28, 594–607.

Ishiguro, K.I., Matsuura, K., Tani, N., Takeda, N., Usuki, S., Yamane, M., Sugimoto, M., Fujimura, S., Hosokawa, M., Chuma, S., et al. (2020). MEIOSIN Directs the Switch from Mitosis to Meiosis in Mammalian Germ Cells. Dev Cell.

Ishishita, S., Matsuda, Y., and Kitada, K. (2014). Genetic evidence suggests that Spata22 is required for the maintenance of Rad51 foci in mammalian meiosis. Sci Rep 4, 6148.

Jensen, R.B., Carreira, A., and Kowalczykowski, S.C. (2010). Purified human BRCA2 stimulates RAD51-mediated recombination. Nature 467, 678–683.

Kojic, M., Yang, H., Kostrub, C.F., Pavletich, N.P., and Holloman, W.K. (2003). The BRCA2-interacting protein DSS1 is vital for DNA repair, recombination, and genome stability in Ustilago maydis. Mol Cell 12, 1043–1049.

Kurzbauer, M.T., Uanschou, C., Chen, D., and Schlogelhofer, P. (2012). The recombinases DMC1 and RAD51 are functionally and spatially separated during meiosis in Arabidopsis. Plant Cell 24, 2058–2070.

La Salle, S., Palmer, K., O’Brien, M., Schimenti, J.C., Eppig, J., and Handel, M.A. (2012). Spata22, a novel vertebrate-specific gene, is required for meiotic progress in mouse germ cells. Biol Reprod 86, 45.

Lam, I., and Keeney, S. (2015). Mechanism and regulation of meiotic recombination initiation. Cold Spring Harb Perspect Biol 7, a016634.

Luo, M., Yang, F., Leu, N.A., Landaiche, J., Handel, M.A., Benavente, R., La Salle, S., and Wang, P.J. (2013). MEIOB exhibits single-stranded DNA-binding and exonuclease activities and is essential for meiotic recombination. Nat Commun 4, 2788.

Mahadevaiah, S.K., Turner, J.M., Baudat, F., Rogakou, E.P., de Boer, P., Blanco-Rodriguez, J., Jasin, M., Keeney, S., Bonner, W.M., and Burgoyne, P.S. (2001). Recombinational DNA double-strand breaks in mice precede synapsis. Nat Genet 27, 271–276.

Marston, N.J., Richards, W.J., Hughes, D., Bertwistle, D., Marshall, C.J., and Ashworth, A. (1999). Interaction between the product of the breast cancer susceptibility gene BRCA2 and DSS1, a protein functionally conserved from yeast to mammals. Mol Cell Biol 19, 4633–4642.

Neale, M.J., and Keeney, S. (2006). Clarifying the mechanics of DNA strand exchange in meiotic recombination. Nature 442, 153–158.

Page, S.L., and Hawley, R.S. (2004). The genetics and molecular biology of the synaptonemal complex. Annu Rev Cell Dev Biol 20, 525–558.

Peters, A.H., Plug, A.W., van Vugt, M.J., and de Boer, P. (1997). A drying-down technique for the spreading of mammalian meiocytes from the male and female germline. Chromosome Res 5, 66–68.

Pispa, J., Palmen, S., Holmberg, C.I., and Jantti, J. (2008). C. elegans dss-1 is functionally conserved and required for oogenesis and larval growth. BMC Dev Biol 8, 51.

Pittman, D.L., Cobb, J., Schimenti, K.J., Wilson, L.A., Cooper, D.M., Brignull, E., Handel, M.A., and Schimenti, J.C. (1998). Meiotic prophase arrest with failure of chromosome synapsis in mice deficient for Dmc1, a germline-specific RecA homolog. Mol Cell 1, 697–705.

Ribeiro, J., Abby, E., Livera, G., and Martini, E. (2016). RPA homologs and ssDNA processing during meiotic recombination. Chromosoma 125, 265–276.

Robert, T., Nore, A., Brun, C., Maffre, C., Crimi, B., Bourbon, H.M., and de Massy, B. (2016). The TopoVIB-Like protein family is required for meiotic DNA double-strand break formation. Science 351, 943–949.

Romanienko, P.J., and Camerini-Otero, R.D. (2000). The mouse Spo11 gene is required for meiotic chromosome synapsis. Mol Cell 6, 975–987.

Royo, H., Prosser, H., Ruzankina, Y., Mahadevaiah, S.K., Cloutier, J.M., Baumann, M., Fukuda, T., Hoog, C., Toth, A., de Rooij, D.G., et al. (2013). ATR acts stage specifically to regulate multiple aspects of mammalian meiotic silencing. Genes Dev 27, 1484–1494.

Scully, R., Chen, J., Plug, A., Xiao, Y., Weaver, D., Feunteun, J., Ashley, T., and Livingston, D.M. (1997). Association of BRCA1 with Rad51 in mitotic and meiotic cells. Cell 88, 265–275.

Sharan, S.K., Pyle, A., Coppola, V., Babus, J., Swaminathan, S., Benedict, J., Swing, D., Martin, B.K., Tessarollo, L., Evans, J.P., et al. (2004). BRCA2 deficiency in mice leads to meiotic impairment and infertility. Development 131, 131–142.

Shinohara, A., and Shinohara, M. (2004). Roles of RecA homologues Rad51 and Dmc1 during meiotic recombination. Cytogenet Genome Res 107, 201–207.

Siaud, N., Dray, E., Gy, I., Gerard, E., Takvorian, N., and Doutriaux, M.P. (2004). Brca2 is involved in meiosis in Arabidopsis thaliana as suggested by its interaction with Dmc1. EMBO J 23, 1392–1401.

Souquet, B., Abby, E., Herve, R., Finsterbusch, F., Tourpin, S., Le Bouffant, R., Duquenne, C., Messiaen, S., Martini, E., Bernardino-Sgherri, J., et al. (2013). MEIOB targets single-strand DNA and is necessary for meiotic recombination. PLoS Genet 9, e1003784.

Tang, Z., Li, C., Kang, B., Gao, G., Li, C., and Zhang, Z. (2017). GEPIA: a web server for cancer and normal gene expression profiling and interactive analyses. Nucleic Acids Res 45, W98–W102.

Vrielynck, N., Chambon, A., Vezon, D., Pereira, L., Chelysheva, L., De Muyt, A., Mezard, C., Mayer, C., and Grelon, M. (2016). A DNA topoisomerase VI-like complex initiates meiotic recombination. Science 351, 939–943.

Wold, S., Boye, E., Slater, S., Kleckner, N., and Skarstad, K. (1998). Effects of purified SeqA protein on oriC-dependent DNA replication in vitro. EMBO J 17, 4158–4165.

Xu, Y., Greenberg, R.A., Schonbrunn, E., and Wang, P.J. (2017). Meiosis-specific proteins MEIOB and SPATA22 cooperatively associate with the single-stranded DNA-binding replication protein A complex and DNA double-strand breaks. Biol Reprod 96, 1096–1104.

Yoshida, K., Kondoh, G., Matsuda, Y., Habu, T., Nishimune, Y., and Morita, T. (1998). The mouse RecA-like gene Dmc1 is required for homologous chromosome synapsis during meiosis. Mol Cell 1, 707–718.

Yoshima, T., Yura, T., and Yanagi, H. (1998). Novel testis-specific protein that interacts with heat shock factor 2. Gene 214, 139–146.

Zhang, J., Fujiwara, Y., Yamamoto, S., and Shibuya, H. (2019). A meiosis-specific BRCA2 binding protein recruits recombinases to DNA double-strand breaks to ensure homologous recombination. Nat Commun 10, 722.

Zhao, W., Vaithiyalingam, S., San Filippo, J., Maranon, D.G., Jimenez-Sainz, J., Fontenay, G.V., Kwon, Y., Leung, S.G., Lu, L., Jensen, R.B., et al. (2015). Promotion of BRCA2-Dependent Homologous Recombination by DSS1 via RPA Targeting and DNA Mimicry. Mol Cell 59, 176–187.

Zickler, D., and Kleckner, N. (2015). Recombination, Pairing, and Synapsis of Homologs during Meiosis. Cold Spring Harb Perspect Biol 7.

Felipe-Medina N., et al. (2020). A missense in HSF2BP causing Primary Ovarian Insufficiency affects meiotic recombination by its novel interactor C19ORF57/MIDAP. bioRxiv, https://doi.org/10.1101/2020.03.05.978007.

